# Pharmacological modulation of dopamine receptors reveals distinct brain-wide networks associated with learning and motivation in non-human primates

**DOI:** 10.1101/2023.12.27.573487

**Authors:** Atsushi Fujimoto, Catherine Elorette, Satoka H. Fujimoto, Lazar Fleysher, Peter H. Rudebeck, Brian E. Russ

## Abstract

The neurotransmitter dopamine (DA) has a multifaceted role in healthy and disordered brains through its action on multiple subtypes of dopaminergic receptors. How modulation of these receptors influences learning and motivation by altering intrinsic brain-wide networks remains unclear. Here we performed parallel behavioral and resting-state functional MRI experiments after administration of two different DA receptor antagonists in macaque monkeys. Systemic administration of SCH-23390 (D1 antagonist) slowed probabilistic learning when subjects had to learn new stimulus-reward associations and diminished functional connectivity (FC) in cortico-cortical and fronto-striatal connections. By contrast, haloperidol (D2 antagonist) improved learning and broadly enhanced FC in cortical connections. Further comparisons between the effect of SCH-23390/haloperidol on behavioral and resting-state FC revealed specific cortical and subcortical networks associated with the cognitive and motivational effects of DA manipulation, respectively. Thus, we reveal distinct brain-wide networks that are associated with the dopaminergic control of learning and motivation via DA receptors.

**Significance Statement:** D1 and D2 receptors are heavily implicated in cognitive and motivational processes, as well as in a number of psychiatric disorders. Despite this, little is known about how selective manipulation of these different receptors impacts cognition through changing activity across brain-wide intrinsic networks. Here, we examined the acute behavioral and brain-wide effects of D1 and D2 receptor-selective antagonists, SCH-23390 and haloperidol, in macaques performing a probabilistic learning task. SCH administration diminished, and haloperidol improved, animals’ task performance. Mirroring these effects on behavior, SCH reduced, and haloperidol increased, the resting-state functional connectivity across brain-wide networks, most notably in the cortico-striatal areas. Thus, our results highlight the opposing effects of D1 and D2 receptor modulation on the brain and behavior.

## Introduction

Dopamine (DA), a neurotransmitter in the central nervous system, plays a critical role in learning, cognitive control, and working memory as well as motivated behavior (Brozoski et al., 1979; Schultz et al., 1997; Volkow et al., 1998; Robbins and Everitt, 2002; Remy and Samson, 2003; Noudoost and Moore, 2011; Ott and Nieder, 2019). DA acts through its binding to various dopamine receptors that are heterogeneously distributed across the brain (Seeman, 1987; Self, 2010). The dopamine D1 and D2 receptors are the most prevalent subtypes of dopamine receptors in both humans and animals and they are heavily implicated in psychiatric conditions such as schizophrenia (Lidow et al., 1998; Brisch et al., 2014).

Extensive research has found that D1 and D2 receptors have distinct roles in learning and motivation. D1 receptor blockade through systemic or local administration in prefrontal cortex disrupts cue-reward association learning and probabilistic reversal learning in rats, while blocking of D2 receptors promotes learning (Eyny and Horvitz, 2003; Zeeb et al., 2009; St Onge et al., 2011; Jenni et al., 2021). Similarly, in macaque monkeys, local administration of a D1 antagonist, SCH-23390, into dorsolateral prefrontal cortex impairs working memory and learning (Sawaguchi and Goldman-Rakic, 1991; Puig and Miller, 2012). By contrast, systemic administration of the D2 antagonist haloperidol, which is widely used to ameliorate positive symptoms of schizophrenia (Settle and Ayd, 1983; Adams et al., 2013), facilitated value discounting (Hori et al., 2021). At the same time, drugs that impact D1 and D2 receptors have differential effects on neural activity. Specifically, earlier PET and SPECT studies reported that the D2 antagonist haloperidol increases cerebral blood flow in healthy individuals and in clinically responsive schizophrenia patients (Buchsbaum et al., 1992; Goldman et al., 1996). Resting-state fMRI studies reported a decrease in the hemodynamic response following administration of D1 antagonist SCH-23390 in rats (Choi et al., 2006), while D2 antagonist haloperidol, or agonist bromocriptine, enhanced dorsal fronto-parietal networks in healthy human subjects (Cole et al., 2013; Vogelsang et al., 2023). Although these studies provided partial evidence as to how D1 and D2 modulation impacts brain-wide intrinsic MRI functional connectivity, how higher doses that are sufficient to robustly modulate behavior would impact brain-wide networks remains unclear.

To address these issues, we conducted parallel behavioral and resting-state functional neuroimaging experiments in macaque monkeys. We found that the selective D1 and D2 receptor antagonists, SCH-23390 and haloperidol respectively (Beaulieu and Gainetdinov, 2011), induced contrasting effects on both behavior and functional connectivity in whole-brain networks. Further, the cortical functional connectivity changes induced by DA antagonists were correlated with task performance, especially when subjects had to learn new stimulus-reward associations. Thus, our results reveal the brain-wide impact of selectively manipulating activity at different DA receptor subtypes, shedding light on the neural networks that are associated with dopamine receptor-dependent cognitive function.

## Materials and Methods

### Subjects

Seven rhesus macaques (*Macaca mulatta,* 7-8 years old, 4 females) served as subjects. All subjects were pair or grouped-housed, were maintained on a 12-h light/dark cycle and had access to food 24 hours a day. During training and testing each subject’s access to water was controlled for 5 days per week. The experiments performed for each subject are summarized in **Table 1**. All procedures were approved by the Icahn School of Medicine Animal Care and Use Committee.

**Table 1.** Assignments of monkeys to resting-state fMRI and behavioral testing conditions. Y and N indicate the condition that the data was collected and not collected, respectively. SCH: SCH-23390 (10 µg/kg), HAL: haloperidol (50 µg/kg). Note that animals assigned to behavioral experiments (Ee, Me, Pi, St) went through all drug treatment conditions.

| ***Subject*** | ***Behavior*** | ***SCH-rsMRI*** | ***HAL-rsMRI*** | ***Saline-rsMRI*** |
| --- | --- | --- | --- | --- |
| ***Ee*** | Y | Y | N | Y |
| ***Me*** | Y | N | Y | N |
| ***Pi*** | Y | Y | Y | Y |
| ***St*** | Y | Y | Y | N |
| ***Bu*** | N | N | N | Y |
| ***Cy*** | N | N | N | Y |
| ***Wo*** | N | N | N | Y |

### Surgery

Prior to training, an MRI compatible head-fixation device (Rogue research, Montréal, Canada) was surgically implanted using dental acrylic (Lang Dental, Wheeling, IL) and ceramic screws (Thomas Research Products, Elgin, IL) in the animals that underwent behavioral testing (monkeys Ee, Me, Pi, St). In a dedicated operating suite using aseptic procedures, anesthesia was induced using ketamine (10 mg/kg, i.m.) and then maintained by isoflurane (2-3%). The skin, fascia, and muscles were opened and retracted. 8-10 MR-compatible ceramic screws were implanted into the cranium and the head fixation device was bonded to the screws using dental acrylic. The muscles, fascia, and skin were then sutured closed. The animals were treated with dexamethasone sodium phosphate (0.4 mg/kg, i.m.) and cefazolin antibiotic (15 mg/kg, i.m.) for one day before and one week after surgery. After surgery and for two additional days, the animals received ketoprofen analgesic (10-15 mg/kg, i.m.); ibuprofen (100 mg) was administered for five additional days and all postoperative medications were given in consultation with veterinary staff. The position of implant was determined based on a pre-acquired T1-weighted MR image.

### Drugs

SCH-23390 hydrochloride (Tocris Bioscience, Minneapolis, MN) and haloperidol (Sigma-Aldrich, St. Louis, MO) were used as our D1 and D2 receptor selective antagonists, respectively. Both SCH and haloperidol were dissolved and diluted in 0.9% saline to achieve the target dose within 1 ml solution. 0.9% saline (1 ml) was also used as a control solution. The solution was prepared fresh on every experimental day using sterile procedures.

### Behavioral experiments

A probabilistic learning task was developed for macaque monkeys (**Fig. 1A**). The task was controlled by NIMH MonkeyLogic software (Hwang et al., 2019) running on MATLAB 2019a (MathWorks, Natick, MA) and presented on a monitor in front of the monkey. In this task, animals were required to choose, using an eye movement, between two visual stimuli presented on either side of a monitor. A trial began with appearance of a fixation spot (white cross) at the center of the screen. The monkey had to acquire and maintain fixation for 1-1.5 sec to initiate a trial. The fixation spot was extinguished, and two visual stimuli were simultaneously presented to the right and left on the screen. The two stimuli presented on each trial were randomly chosen from a larger pool of three visual stimuli that were associated with different reward probabilities (0.9, 0.5, and 0.3) (**Fig. 1B**). Each trial therefore fell into three categories based on the reward probabilities of the options presented: High-Low (0.9-0.3), High-Mid (0.9-0.5), and Mid-Low (0.5-0.3). Stimuli were either novel at the beginning of each block of 100 trials (novel block) or subjects had previously learned about the reward-probability associated with each image and were highly familiar with them (familiar block). Once stimuli were presented, subjects were required to move their eyes toward either right or left stimulus option (‘response’) within 2 seconds. Following a response, the chosen stimulus remained on screen for 0.3 sec and then was removed, and a fluid reward was then immediately delivered based on the probability of the chosen option. Subsequently an inter-trial interval (ITI, 3-3.5 sec) followed. A trial with a fixation break during the fixation period or with no response within the response window was aborted; all stimuli were extinguished immediately, and an ITI started. The same trial was repeated following an aborted trial.

**Figure 1.**
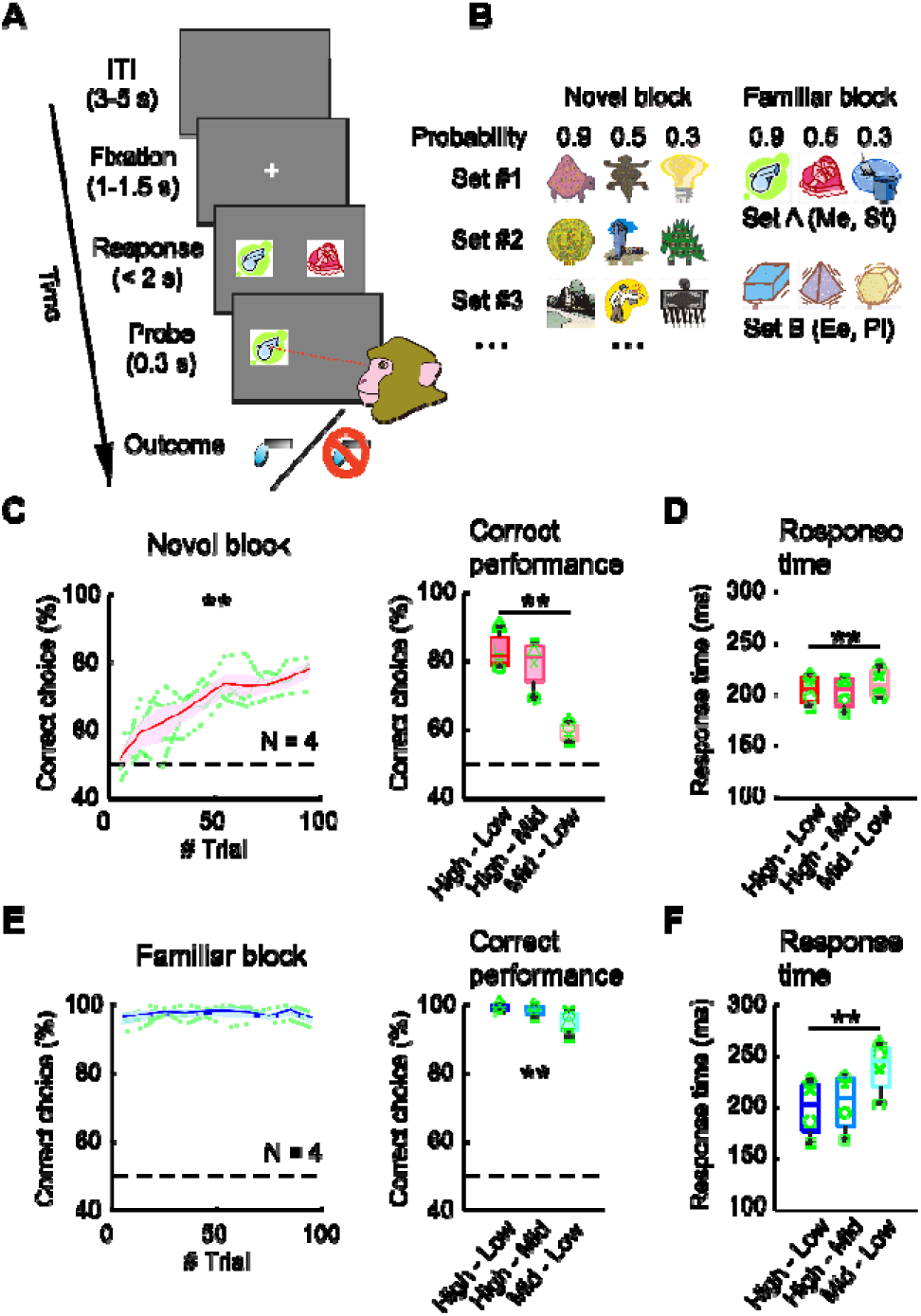
Behavioral task and baseline behavioral performance. (**A**) Trial sequence. Animals were required to respond to one of two visual stimuli on the screen by eye movement to acquire a drop of juice. (**B**) Stimulus sets. Stimuli were pictures that were associated with different reward probabilities (0.9, 0.5, 0.3). In novel blocks (left), a new set of three pictures was used in each block. In familiar blocks (right), a fixed set of pictures was prepared for each monkey and used repeatedly throughout the experiment. (**C**) Task performance in novel blocks. Correct performance was gradually increased over trials in a block (left) and depending on which stimuli were paired in the trial (right). Dashed lines indicate chance level, and the plots show mean and standard error. Green lines indicate the average performance of each animal. Box plots indicate the median, 25^th^ and 75^th^ percentiles, and the extent of data points obtained in the 2^nd^ half of each block. High-Low: 0.9-0.3, High-Mid: 0.9-0.5, Mid-Low: 0.5-0.3. Symbols indicate individual animals. (**D**) Response time (RT) in novel blocks reflected reward probability of the stimulus pair. (**E and F**) Behaviors in familiar blocks. Correct performance was stable throughout the block. Performance and RT reflected reward probability. Conventions are the same as **C-D**. ** p < 0.01, interaction of trial bin by block type, 2-way repeated-measures ANOVA, or main effect of stimulus pair, 1-way repeated-measures ANOVA.

The animals performed 4-6 blocks in which the novel or familiar stimuli were pseudorandomly interleaved in hour-long sessions. The monkeys were trained for 3-6 months before behavioral experiments with drug injections or resting-state fMRI scans. The I.M. injection of saline, SCH-23390 (10, 30, or 50 µg/kg), or haloperidol solution (5 or 10 µg/kg) was performed 15 minutes prior to the task start. Each monkey completed at least 3 sessions at each dose level for each drug for a total of 80-138 total blocks per monkey. The order of treatment was randomized, and injections were at least a day (SCH-23390) or week apart (haloperidol) to avoid potential prolonged effects of the drug, in accordance with known pharmacokinetics of the drugs in macaque monkeys (Hori et al., 2021).

### Resting-state fMRI data acquisition

The scans were performed under the same protocol we previously developed for macaque monkeys (Fujimoto et al., 2022; Elorette et al., 2024). In brief, following sedation with ketamine (5mg/kg) and dexmedetomidine (0.0125mg/kg) the animals were intubated. They were then administered (i.v.) monocrystalline iron oxide nanoparticle or MION (10 mg/kg, BioPAL, Worcester, MA), and three EPI functional scans (1.6 mm isotropic, TR/TE 2120/16 ms, flip angle 45°, 300 volumes per each run) were obtained, along with a T1-weighted structural scan (0.5 mm isotropic, MPRage TR/TI/TE 2500/1200/3.27 ms, flip angle 8°) (pre-injection scans). Following drug i.v. injection (saline, SCH-23390, or haloperidol) and 15 minutes waiting period, another set of three functional scans was acquired (post-injection scans). Low-level isoflurane (0.7-0.9%) was used to maintain sedation through a session so that neural activity was preserved while minimizing motion artifacts. Vital signs (end-tidal CO_2_, body temperature, blood pressure, capnograph) were continuously monitored and maintained as steadily as possible throughout an experimental session. The doses of drugs used in the scans (50 µg/kg and 10 µg/kg for SCH and haloperidol, respectively) were pre-determined based on a prior PET study to achieve up to 70-80% occupancy of the DA receptors in macaques (Hori et al., 2021).

### Behavioral data analyses

All behavioral data was analyzed using MATLAB 2019a. Choice performance was defined as the proportion of trials in a block (100 trials) in which monkeys chose an option associated with higher reward probability in the stimulus pair presented. Response time (RT) was defined as the duration from the timing of visual stimuli presentation to the timing of response initiation. Choice performance was computed for bins of 10 trials at each block and averaged for each subject, then finally averaged across subjects for each block type. We reasoned that a significant interaction (p < 0.05) of trial bin by block type with 2-way repeated measures ANOVA (trial bin: 1-10 × block type: novel or familiar) indicated that there was an improvement in performance due to successful learning in novel blocks but not in familiar blocks. Choice performance and RT on each stimulus pair in the latter half of each block were assessed by 1-way repeated-measures ANOVA (stimulus pair: 0.9-0.3, 0.9-0.5, 0.5-0.3) for each block type in saline sessions. The effect of SCH-23390 or haloperidol injection on choice performance and RT was assessed by 3-way repeated-measures ANOVA (block type: novel or familiar × stimulus pair: 0.9-0.3, 0.9-0.5, 0.5-0.3 × drug dose: 0, 10, 30, 50 µg/kg SCH-23390, and 0, 5, or 10 µg/kg haloperidol). To further assess the effects of drugs on each block type, we also performed 2-way repeated-measures ANOVA (stimulus pair: 0.9-0.3, 0.9-0.5, 0.5-0.3 × drug dose: 0, 10, 30, 50 µg/kg for SCH-23390, and 0, 5, or 10 µg/kg for haloperidol, respectively). All multi-way ANOVA was performed by using MATLAB built-in function *anovan* with monkeys modeled as a random effect.

We also performed a model fitting analysis for the choice data in novel blocks employing a standard reinforcement learning model with a softmax choice function (Sutton and Barto, 1981; Rudebeck et al., 2017b) described as below:

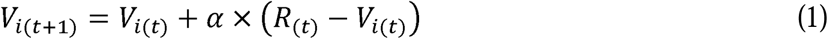

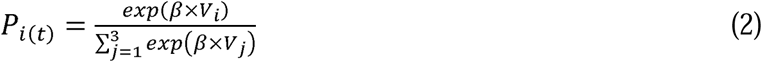

Where α and β represent learning rate and inverse temperature, respectively. *V_i(t)_* and *R_(t)_*indicate the value of the chosen option *i* and outcome on trial *t*. *P_i(t)_* indicates the choice probability of option *i* on trial *t*. Then the log-likelihood (LL) and the Bayesian Information Criterion (BIC) were calculated for each block to assess how well the model fitted the data:

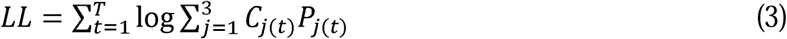

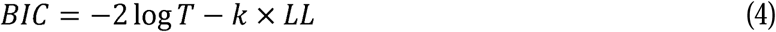

Where *T* and *k* denote the size of trial block and the number of parameters, respectively. *C_j(t)_* = 1 when the subject chooses option *j* in trial *t*, and *C_j(t)_* = 0 for all unchosen options. The learning rate and inverse temperature were estimated using MATLAB function *fminsearchbnd* to select parameters by minimizing the log-likelihood function for each block. The best-fit parameters were averaged for each drug condition, and the dose-dependent effects of drugs as well as BIC were assessed by 1-way repeated-measures ANOVA (drug dose: 0, 10, 30, 50 μg/kg for SCH-23390, and 0, 5, or 10 μg/kg for haloperidol, respectively).

### fMRI data analysis

The detail of preprocessing steps for functional imaging data was described in our previous study (Fujimoto et al., 2022). In brief, all functional imaging data was initially converted to NIFTI format and preprocessed with custom AFNI/SUMA pipelines (Cox, 1996; Jung et al., 2021; Fujimoto et al., 2022). The T1 weighted image from each session was skullstripped (Wang et al., 2021) and then warped to the standard NMT atlas space (Seidlitz et al., 2018). The EPI data were further preprocessed using a customized version of the AFNI NHP preprocessing pipeline (Jung et al., 2021). The first 3 TRs of each EPI were removed to eliminate any magnetization effects. Then, the images went through slice timing correction, motion correction, alignment to T1w image, warping to standard space, blurring, and then converted to percent signal change. Finally, motion derivatives from each scan along with CSF and WM signals were regressed and the residuals of this analysis were used in the following analysis.

The functional connectivity (FC) analysis was performed using 3dNetCorr function in AFNI (Cox, 1996; Taylor and Saad, 2013). The regions of interest (ROIs) were defined based on the cortical hierarchical atlas (CHARM) (Jung et al., 2021) and subcortical hierarchical atlas (SARM) (Hartig et al., 2021) for rhesus macaques, both at level 4. The matrices of FC across all ROI pairs, or connectomes, were Fisher’s z-transformed for each session, and the pre-injection connectome was subtracted from post-injection connectome. Then, the connectomes representing the drug-induced change in FC (ΔFC) were averaged within treatment conditions (SCH-23390, haloperidol, saline). To statistically determine the effects, the ΔFCs derived from each ROI were averaged and compared to a null distribution (α = 0.05 with Bonferroni’s correction, rank-sum test). The connectomes were also visualized in the circular plot with the threshold set at z = 0.1 (absolute value) created using the circularGraph toolbox run in MATLAB (Kassebaum, 2023). Separately, we also analyzed the whole-brain FC using a dorsal and ventral striatum seed. Correction for multiple comparisons was performed using 3dClustSim, which computed the cluster-size threshold based on 10000 iteration of Monte Carlo simulations in AFNI (Cox, 1996). The combination of initial thresholding at p < 0.01 and the cluster-size threshold at 6 voxels corresponds to corrected p < 0.05.

The relationship between the connectome and behavioral data (correct performance and RT) and between the connectome and RL parameters (learning rate and inverse temperature) were analyzed on the data where ΔFC and behavioral data were obtained under the same drug condition, and all drug conditions (saline, SCH, haloperidol) were combined. The correlation analysis was performed separately for each functional connection or ROI pair, and a matrix of correlation coefficients (R) was created. A permutation test was performed for each functional connection by comparing R^2^ computed from real data and that derived from shuffled data with randomized behavioral sessions 1000 times. The correlation matrix was also projected into a brain map of macaque monkeys by connecting the center of each ROI with a line reflecting the R-value and sign (positive or negative) of correlation as the line width and color, respectively. For visualization purposes the fraction of connections that showed strong behavior-ΔFC correlation (top 5%) were plotted. The R values in the matrix were averaged across functional connections for each of cortico-cortical, cortico-subcortical, and subcortico-subcortical ROI pairs and compared to the null distribution (rank-sum test).

## Results

### Distinct effects of dopamine receptor antagonists on probabilistic stimulus-reward learning

Four macaque monkeys were trained to perform a probabilistic learning task for fluid rewards. On each trial, the animals were free to choose between the two visual stimuli by making an eye movement to obtain a juice reward (**Fig. 1A**). The stimuli presented on each trial were randomly chosen from a set of three stimuli that were associated with distinct reward probabilities (0.9, 0.5, and 0.3) (**Fig. 1B**). Subjects completed 100-trial blocks with either stimuli that were novel at the start of each block (novel blocks) or that they had previously learned (familiar blocks).

In novel blocks with saline administration, monkeys gradually learned to discriminate between the different stimuli (**Fig. 1C**). By contrast, in the familiar blocks subjects reliably maintained a high and stable performance throughout a given block, suggesting memory-guided choices (**Fig. 1E**). A two-way repeated-measures ANOVA (trial bin: 1-10 × block type: novel or familiar) revealed a significant interaction of trial bin by block type on choice performance (p < 0.01, F_(9,1097)_ = 8.0), confirming the difference between novel and familiar blocks. Subjects’ choice performance was also influenced by which stimuli were presented as options on each trial. A one-way repeated-measures ANOVA (stimulus pair: 0.9-0.3, 0.9-0.5, 0.5-0.3) revealed a significant main effect of stimulus pair on choice performance in both novel and familiar blocks (novel blocks: p < 0.01, F_(2,168)_ = 16.3; familiar blocks: p < 0.01, F_(2,153)_

= 15.3, **Fig. 1C and E**). Additionally, response time (RT) reflected the reward probability of available options in both block types, such that RT was shorter for trials in which the high reward probability stimulus was presented (one-way repeated-measures ANOVA, novel blocks: p = 0.027, F_(2,168)_ = 3.7; familiar blocks: p < 0.01, F_(2,153)_ = 53.8, main effect of stimulus pair, **Fig. 1D and F**). Importantly, the patterns of behavior were consistent across all subjects in both the novel and familiar blocks (**Fig. 1C-F**).

Following administration of dopamine receptor antagonists, behavioral performance was impacted (**Fig. 2A and B**). A set of larger ANOVA models including both SCH-23390 and haloperidol conditions (drug × block type × stimulus pair) revealed a significant interaction of drug by block type (p < 0.01, F_(5,1110)_ = 3.6), indicating that dopamine receptor antagonists specifically impact performance when monkeys have to learn novel stimulus-reward associations. Notably, we found that SCH-23390 tended to decrease subjects’ performance in novel blocks (p = 0.061, F_(3,441)_ = 2.5, main effect of drug dose, 2-way repeated-measures ANOVA), while it did not affect the performance in blocks with familiar stimuli (p = 0.90, F_(3,399)_ = 0.20) (**Fig. 2C**). The treatment also affected RT such that higher doses of SCH increased RT in both novel and familiar blocks (novel blocks: p < 0.01, F_(3,441)_ = 24.0; familiar blocks: p = 0.015, F_(3,399)_ = 3.5) (**Fig. 2D**). In contrast to SCH-23390, haloperidol increased subjects’ correct performance in novel blocks (p = 0.037, F_(2,342)_ = 3.3), while it did not affect the performance in familiar blocks (p = 0.63, F_(2,309)_ = 0.46) (**Fig. 2E**). Notably, administration of haloperidol did not affect subjects RTs in either novel or familiar blocks (p > 0.53), suggesting negligible effects on monkeys’ motivation at the range of doses we used (**Fig. 2F**). Thus, dopamine receptor antagonists induced opposing effects on learning novel probabilistic stimulus-reward associations at the higher doses that we used, while they had no discernable impact on familiar associations.

**Figure 2.**
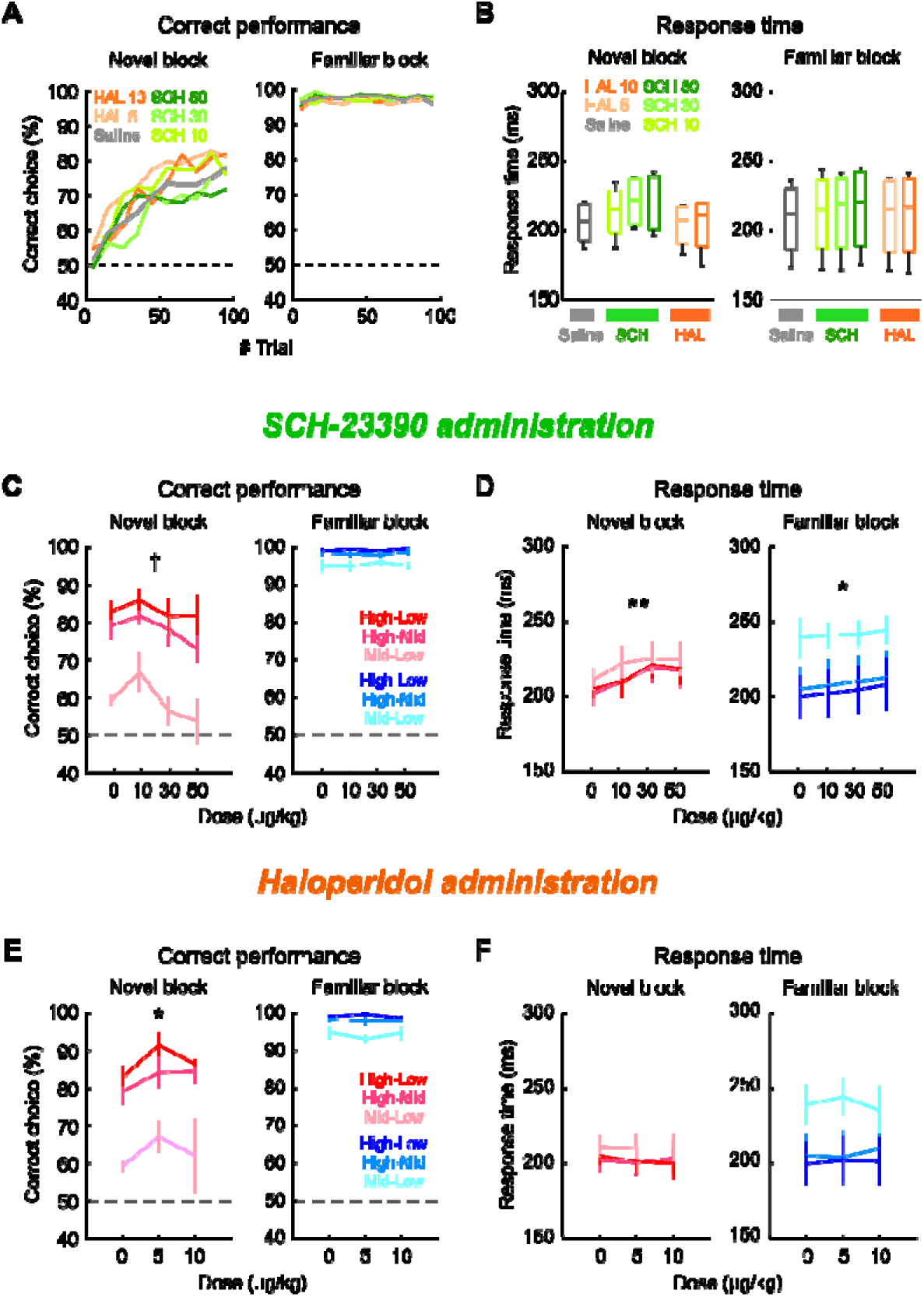
Effects of DA receptor antagonists on behaviors. (**A-B**) Overall summary of drug effects on behaviors. (**A**) Averaged performance (proportion of correct choice) plotted against the trial number for novel (left) and familiar (right) blocks, respectively. Line colors indicate the drug type and shade indicates the dose, orange shades (Haloperidol), green shades (SCH-23390), grey (Saline). (**B**) Drug effects on response time (RT). Box plots indicate the median, 25th and 75th percentiles, and the extent of data points. (**C-F**) Drug effects collapsed by drug dose and stimulus pair. (**C**) Task performance in SCH-23390 sessions. Correct performance tended to decrease when higher dose of SCH was administered in novel blocks (left) but did not change in familiar blocks (right). The colors of lines indicate stimulus pairs. (**D**) RT in SCH-23390 sessions. RT increased following SCH injection. (**E and F**) Haloperidol sessions. Conventions are the same as **C-D**. †p < 0.10, *p < 0.05, **p < 0.01, 2-way repeated-measures ANOVA. Symbols indicate individual animals.

We also assessed the effect of drugs during learning by a model fitting analysis employing a standard two parameter reinforcement learning model (see Materials and Methods). The model was fitted to the animals’ choice data in each block of the novel condition (**Fig. 3A**), and the average of best-fit parameters were computed for each drug condition (**Fig. 3B-C**). This analysis revealed that haloperidol, but not SCH-23390 administration, tended to decrease inverse temperature (haloperidol: p = 0.073, F_(2,110)_ = 2.7, SCH-23390: p = 0.81, F_(3,141)_ = 0.32, main effect of drug dose with 1-way repeated-measures ANOVA), while neither drug changed the animals’ learning rate (p > 0.20). Importantly, we did not find a significant difference between the model fits as measured by the Bayesian Information Criteria (BIC), across the different levels of SCH-23390 or haloperidol (p > 0.25, main effect of drug dose with 1-way repeated-measures ANOVA). This result indicates that D2 receptor manipulation impacted the animals’ degree of exploration, while D1 receptor antagonism did not affect either process, during learning.

**Figure 3:**
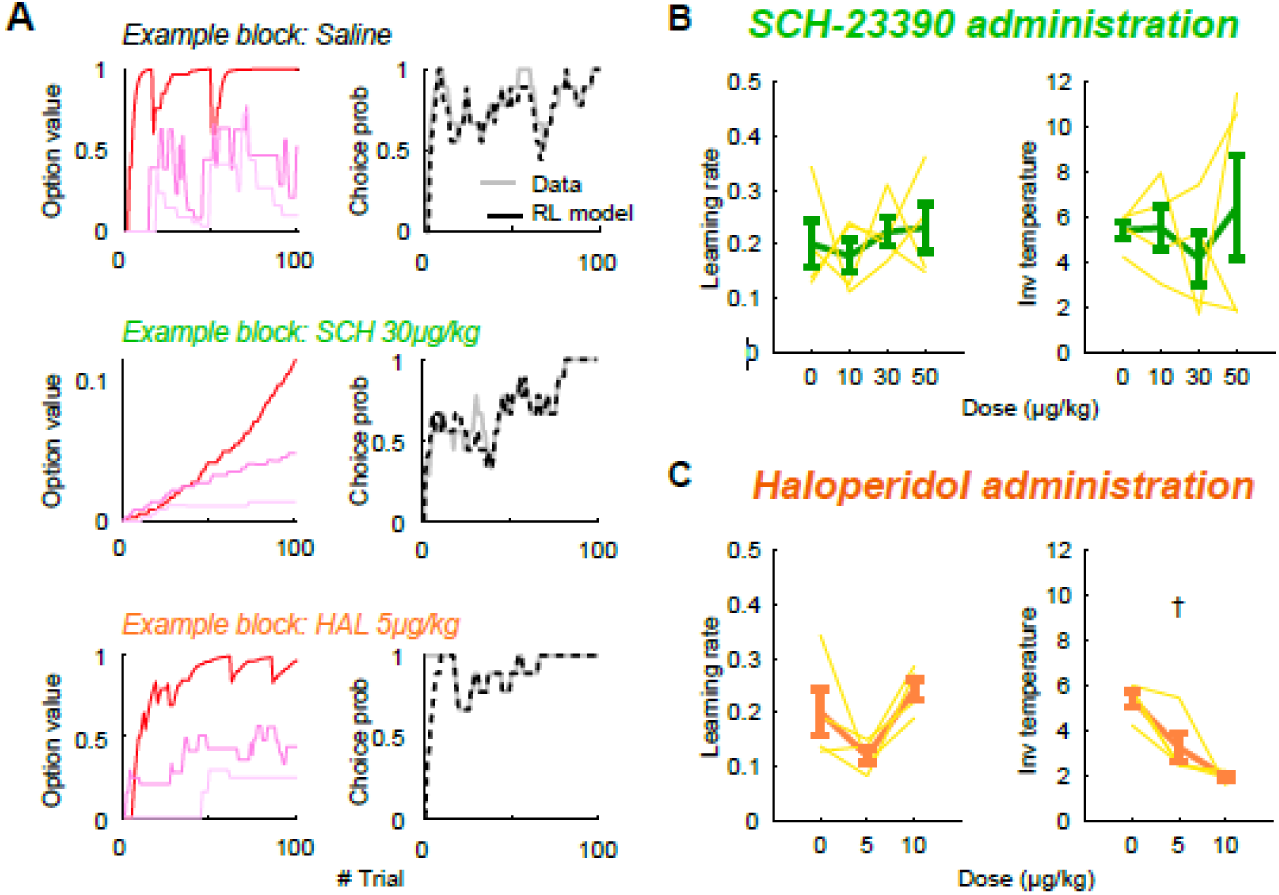
Reinforcement learning model fitting. (**A**) RL model fitting on choice data in an example block with administration of saline (top), SCH-23390 (middle), and haloperidol (bottom). The left panels show the transition of the model estimated value in example blocks (line colors indicate stimuli). The right panels show the animal’s choice probability (gray solid lines) and the estimated choice probability based on RL model (black broken lines) in the same blocks. (**B**) The dose-dependent effects of SCH-23390 on learning rate (left) and inverse temperature (right). Thin yellow lines indicate the data from individual animals. (**C**) The dose-dependent effects of haloperidol on learning rate (left) inverse temperature (right). †p < 0.10.

### Contrasting effects of dopamine receptor antagonists on fronto-striatal functional connectivity

Given the clear differences between D1 and D2 receptor antagonism on monkeys’ performance of the probabilistic task, we next set out to determine which networks might be most influenced by our two DA receptor antagonists and therefore potentially driving the behavioral effects. To do this we analyzed resting-state functional images that were obtained in parallel to the behavioral experiments. In addition to the cohort that completed behavioral testing detailed above, three other macaques also underwent saline scans to serve as additional baseline data for our analyses (see **Table 1**).

First, to assess the effects of the drugs on an area known to be high in D1 and D2 receptors that has also been implicated in associative learning (Balleine et al., 2007; Clarke et al., 2008; Vo et al., 2014; White and Monosov, 2016), we analyzed the change in dorsal striatum functional connectivity (FC) induced by administration of either SCH-233980 or haloperidol. During baseline imaging, before the injection of either drug, signal in the dorsal striatum ROI (SARM atlas) (Hartig et al., 2021) exhibited high levels of correlation with frontal cortex, including parts of ventrolateral prefrontal cortex (vlPFC) and orbitofrontal cortex (OFC) (**Fig. 4A**). As expected, injection of saline had little effect on dorsal striatum FC with the rest of the brain (**Fig. 4B**). By contrast, administration of SCH-23390 induced broad changes in dorsal striatum FC (**Fig. 4C**). Notably, D1-receptor antagonism specifically decreased dorsal striatum FC with OFC and lateral prefrontal cortex, while increasing correlations within the dorsal striatum itself (p < 0.05, cluster-level correction). By contrast, administration of haloperidol significantly increased FC in frontal-striatal circuits, most notably between striatum and parts of the medial OFC and vlPFC (p < 0.05, **Fig. 4D**), while showing minimal change in FC within the dorsal striatum. We also analyzed the drug effects on the whole-brain FC using the ventral striatum as the seed ROI (SARM level 4 atlas), as this part of the striatum is also implicated in reinforcement learning (van der Meer and Redish, 2011; Averbeck and Costa, 2017) (**Fig. 4E-H**). We found that SCH-23390 and haloperidol induced FC changes similar to those we observed in dorsal striatum, although both the baseline FC and the effects of the drugs were relatively small and there were no significant drug-induced changes in connectivity with frontal cortex (p > 0.05, cluster-level correction). Thus, D1 and D2 receptor antagonism appears to have opposing effects on dorsal striatum FC in macaques, especially with the parts of frontal cortex involved in probabilistic learning (Rudebeck et al., 2017a; Murray and Rudebeck, 2018).

**Figure 4.**
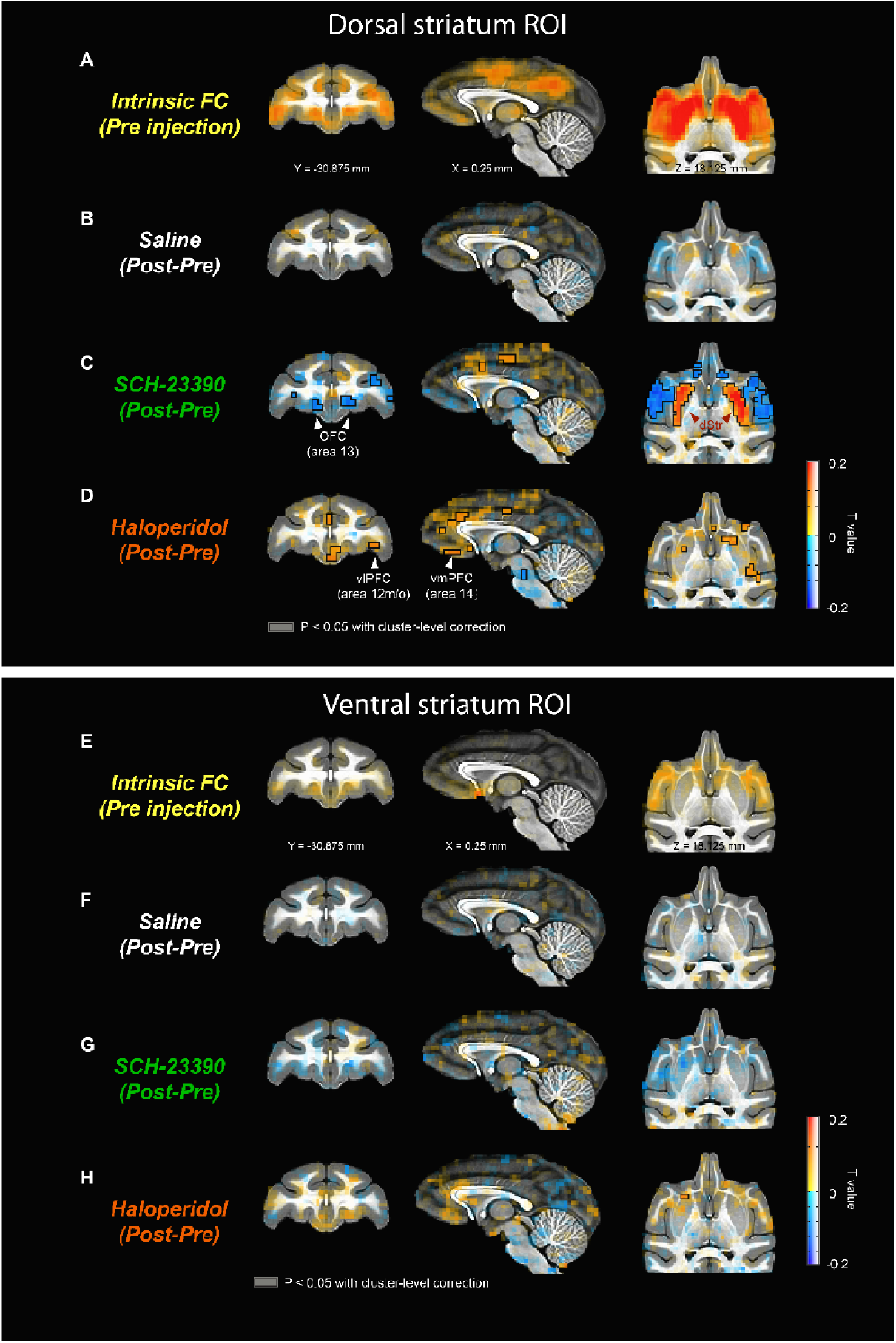
Functional connectivity analysis. (**A**) Whole-brain FC computed using dorsal striatum as ROI to pre-injection images. Coronal (left), sagittal (middle), and axial planes (right) are shown. Colors indicate strength of FC (T-value). (**B**) Changes in dorsal striatum FC from pre-to post-saline injection scans. (**C**) SCH-23390 effects on FC. (**D**) Haloperidol effects on FC. The voxels enclosed in black lines are the clusters with a significant change in dorsal striatum FC (p < 0.05, cluster-level correction). Note that the statistical tests were performed only for subtraction images in **B-D**. dStr: dorsal striatum, OFC: orbitofrontal cortex, vlPFC: ventrolateral prefrontal cortex, vmPFC: ventromedial prefrontal cortex. (**E-H**) Whole-brain FC changes computed using ventral striatum as ROI. Conventions are the same as in **A-D**.

### Functional connectome analysis reveals distinct network signatures associated with dopamine receptor antagonism

To further characterize the impact of the D1 and D2 receptor antagonists on brain-wide networks, we performed atlas-based full connectome analyses. Here we used pre-determined anatomical ROIs from the cortical and subcortical atlas of the macaque monkey (CHARM and SARM atlas, respectively) (Hartig et al., 2021; Jung et al., 2021) and normalized FCs (z-value) were computed for all ROI pairs to produce connectomes. The pre-injection connectomes were similar to those reported previously (Grayson et al., 2016; Fujimoto et al., 2022) (**Fig. 5A, left column**). As expected, injections of saline were not associated with systematic changes in FCs (ΔFCs) of the cortical and subcortical connectome (**Fig. 5A, top row**). By contrast, SCH-23390 injection induced an overall decrease in FCs primarily between cortical regions (**Fig. 5A, middle row**), whereas haloperidol injection induced the opposite pattern of effects on FCs (**Fig. 5A, bottom row**). Indeed, the average z-value for each pair of ROIs showed contrasting effects overall, where SCH-23390 decreased and haloperidol increased FC between cortical sites (p < 0.05 with Bonferroni correction, rank-sum test, **Fig. 5B**).

**Figure 5.**
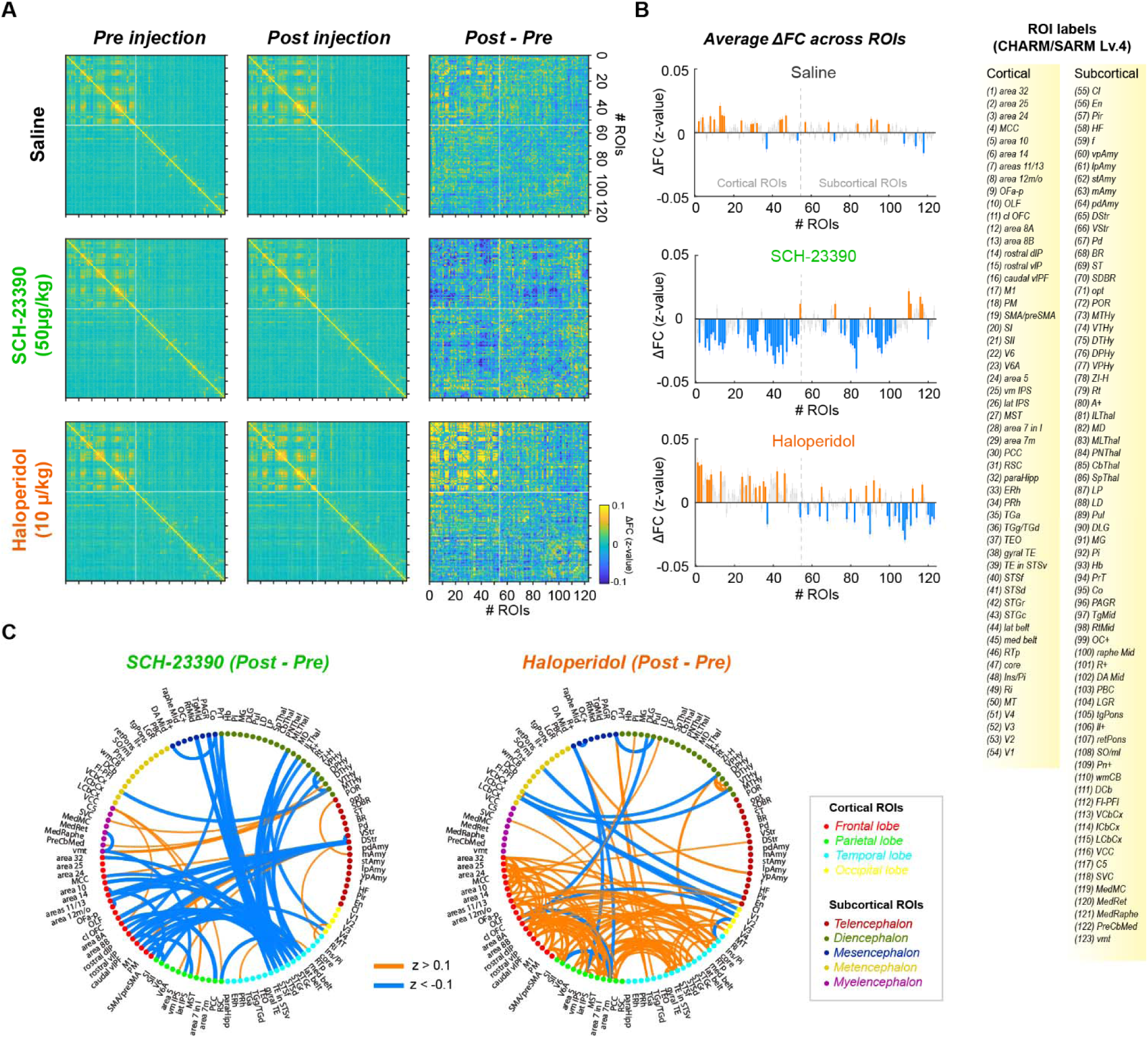
Connectome analysis. (**A**) Connectome for pre-injection (left), post-injection (middle), and their difference (ΔFC, right) are shown for sessions with injection of saline (top row), SCH-23390 (middle row), and haloperidol (bottom row), respectively. X and Y axes are the number of ROIs defined by macaque brain hierarchical atlas (CHARM/SARM atlas at level 4). Colors indicate the FC of each pair of ROIs (z-value). White lines on connectome divide cortical and subcortical ROIs. (**B**) Bar plots showing average ΔFC for saline (top), SCH-23390 (middle), and haloperidol (bottom) sessions. ΔFC (z-value) is averaged for each ROI. Orange and blue bars indicate significant ΔFC from zero (p < 0.05, Bonferroni correction). Dashed lines divide cortical and subcortical ROIs. ROI labels from CHARM/SARM level 4 are shown on the right. (**C**) Circular plots depicting the effects of injection of SCH-23390 (left) or haloperidol (right) on the whole-brain FC. Seed region labels correspond to ROI labels in **B**. The changes in FC are indicated by color (orange: positive changes, blue: negative changes) and width of lines (absolute z-value changes > 0.1). The color of each seed indicates the region defined by CHARM/SARM level 1 (inset).

We further visualized the changes in FC following injections of SCH-23390 or haloperidol by projecting the ΔFC connectomes onto circular plots (absolute difference in z-value > 0.1, **Fig. 5C**). This approach revealed unique patterns of network-level effects induced by SCH-23390 and haloperidol. Specifically, SCH-23390 was associated with a general decrease in cortico-cortical FC in frontal and temporal areas, fronto-striatal FC, and meso/thalamo-cortical FCs. In contrast, haloperidol primarily caused an increase in cortico-cortical FC in frontal, parietal, and temporal areas as well as fronto-striatal FC. In addition to increased FCs, some connections such as midbrain to parietal cortex FC were decreased by treatment with haloperidol. Overall, mirroring our earlier behavioral analyses, SCH-23390 and haloperidol induced contrasting effects in brain-wide FCs, and in particular induced opposite effects in fronto-striatal and cortico-cortical FCs.

### Network correlates of behavioral performance associated with dopaminergic function

The prior analysis shows that the behavioral effects of D1 and D2 receptor antagonism are associated with distinct changes in brain-wide FC. To directly compare behavioral and neuroimaging datasets, we next examined whether the pharmacologically induced changes in resting-state functional connectivity (ΔFC) are related to the effects on behavioral data, either correct performance or RT, that were obtained after the administration of matching doses of the same D1- and D2-antagonists. This allowed us to assess whether changes in FC were related to changes in behavioral responses during a task, even though they were tested under different settings.

We first chose several areas known to be involved in probabilistic learning, namely OFC, vlPFC, dorsal and ventral striatum, mediodorsal thalamus, and midbrain (Clarke et al., 2008; Rudebeck et al., 2017a; Murray and Rudebeck, 2018), and specifically analyzed functional connectivity between those structures. Notably, we found that dorsal striatum-to-OFC ΔFC was significantly correlated with the correct performance in novel blocks (p < 0.01, r = 0.32) (**Fig. 6A and B**), while there was no association between performance in the familiar blocks (p = 0.96) or RTs (p > 0.38) (**Fig. 6C**). The same pattern was seen between OFC-to-12m/o (rostral vlPFC) ΔFC and behavior where a positive correlation was observed between the FC changes and the performance in the novel block (p < 0.01, r = 0.34) (**Fig. 6D-F**). This result indicates that these connections may be involved specifically in learning rather than in the probabilistic choice in general. By contrast, we found a distinct effect on connectivity between mediodorsal thalamus and caudal vlPFC (area 12o): ΔFC between these structures showed no significant correlation with the animals’ performance during the novel blocks (p = 0.79, r = 0.03), but there was a significant negative correlation with performance during the familiar blocks (p = 0.016, r = -0.30) (**Fig. 6G and H**). However, ΔFC between these regions was related to subject’s RTs in both conditions (novel blocks: p < 0.01, r = -0.45; familiar blocks: p < 0.01, p = -0.41) (**Fig. 6I**). Similarly, ventral striatum-to-midbrain ΔFC was also related to RT effects in both conditions (p < 0.032) and showed no significant association with correct performance (p > 0.074) (**Fig. 6J-L**). The strong negative correlation observed between ΔFC and the animals’ RTs suggests that these connections are involved in functions such as motivation or motor control.

**Figure 6.**
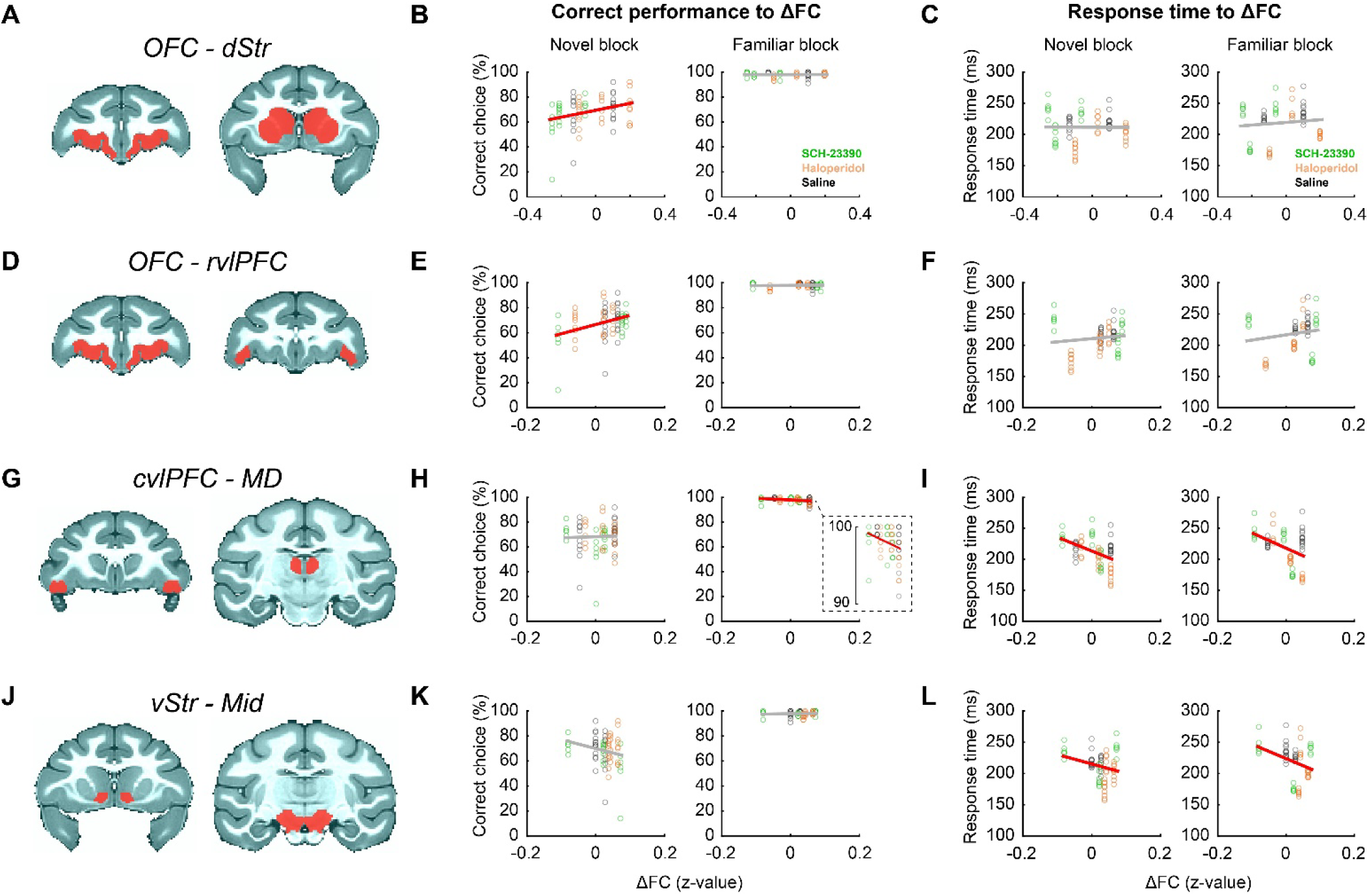
Direct comparison of behaviors and resting-state FC. (**A-C**) Correlation between orbitofrontal cortex to dorsal striatum ΔFC and task performance in novel (left) and familiar (right) blocks (**B**) or response time (**C**). Red areas in brain map show bilateral ROIs. Plots indicate behavioral data and corresponding ΔFC in saline (black), SCH-23390 (green), and haloperidol (orange) sessions. Red and gray lines on scatter plots indicate significant (p < 0.05, linear regression analysis) and non-significant relationships between behavior and ΔFC, respectively. (**D-F**) Correlation between orbitofrontal cortex to rostral part of ventrolateral prefrontal cortex ΔFC and behaviors. (**G-I**) Correlation between the caudal part of ventrolateral prefrontal cortex to mediodorsal thalamus ΔFC and behaviors. Inset is a magnification image for the correlation between familiar block performance and ΔFC. (**J-L**) Correlation between ventral striatum to midbrain ΔFC and behaviors. Conventions are the same as **A-C**.

Next, we extended the approach described above on the full connectome of all ROI pairs and measures of behavior (**Fig. 7**). **Figure 7A** depicts the functional connections where we observed a strong correlation between ΔFC and task performance. The brain map indicates that the task performance in novel blocks was positively correlated to cortico-cortical and cortico-subcortical ΔFCs (**Fig. 7A, left**). Interestingly, the pattern was strikingly different when we analyzed the familiar block; strong correlations were observed mainly in subcortical regions, while cortico-cortical ΔFCs were less correlated to the performance (**Fig. 7A, right**). The full correlation matrix further revealed the detail of these differences (**Fig. 7C**). Notably, there was a strong correlation between correct performance and ΔFCs in cortical areas including frontal, parietal, and temporal regions, as well as in these regions’ functional connections to striatum in the novel blocks (**Fig. 7C, left**). A permutation test with shuffled behavioral sessions (1000 iterations) confirmed that the correlations in those functional connections were significantly greater than the chance (> 95% confidence interval, **Fig. 7E**).

**Figure 7.**
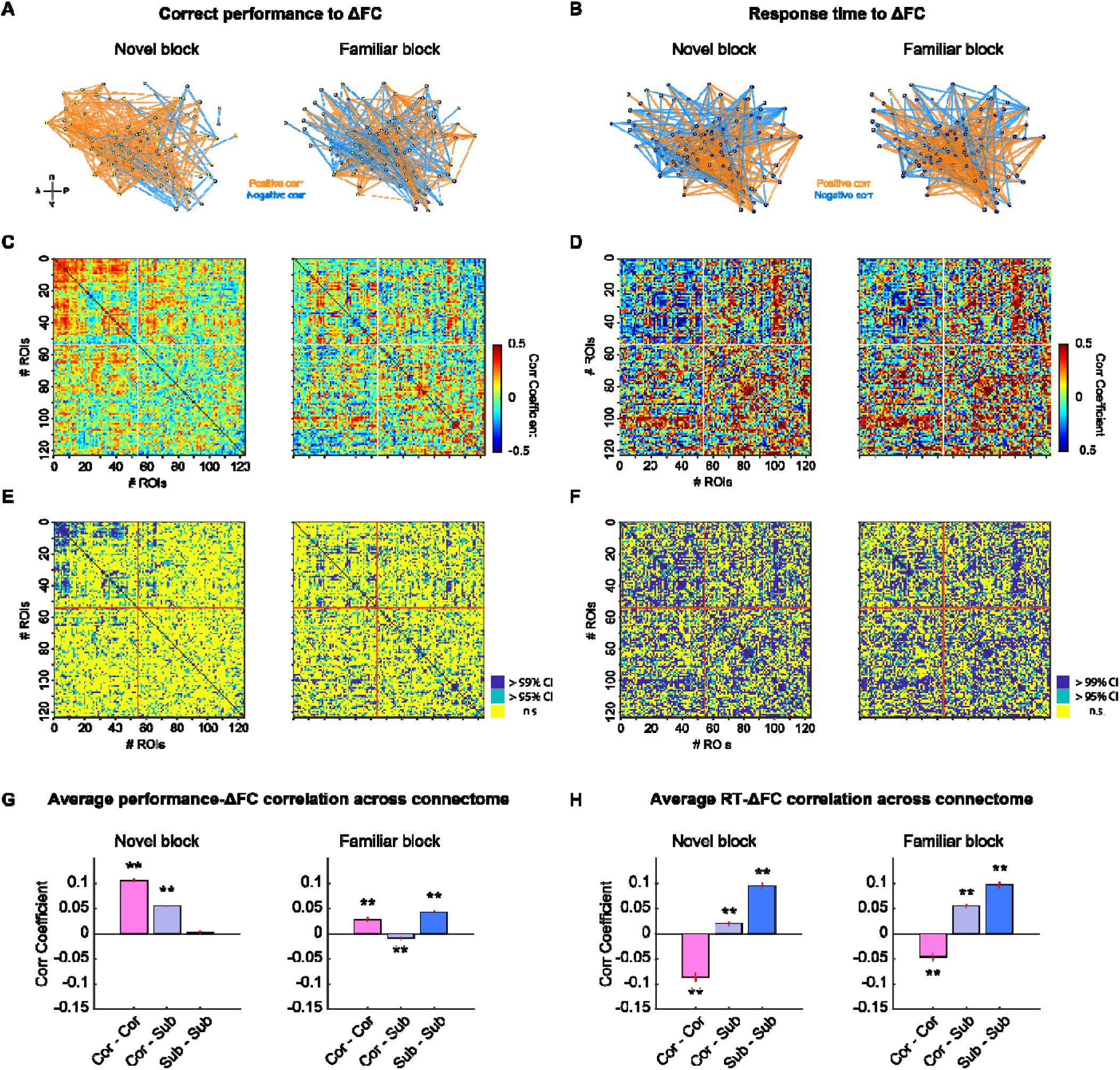
Whole-brain network correlation to behaviors across all drug conditions. (**A**) Strength of performance-ΔFC correlation projected into brain map. The top 5% of connections that showed strong behavior-ΔFC correlation are visualized. Black dots indicate the center of mass of ROIs. The strength and direction of changes in FC are depicted as the width and color (orange: positive, blue: negativ ) of lines, respectively. (**B**) Strength of RT-ΔFC correlation projected into brain map. (**C**) Correlation matrix depicting relationship between changes in FC and task performance in novel (left) and familiar (right) blocks. The ROIs used are the same as in Fig. 5A. Colors indicate performance-ΔFC correlation coefficient. (**D**) Correlation between changes in FC and RT. (**E and F**) Functional connections (ROI pairs) that showed a significant correlation between FC changes and correct performance (**E**) or RT (**F**) (> 95% CI, permutation test) are shown in the matrix used in **C-D**. (**G**) Bar plots depicting average performance-ΔFC correlation coefficients calculated for cortico-cortical, cortico-subcortical, and subcortico-subcortical connections separately, in novel (left panel) and familiar (right panel) blocks. (**H**) Averaged correlation coefficients for RT to ΔFC. **p < 0.01, rank-sum test.

In the familiar blocks, the correlations between cortical areas and performance were less strong, although the change in some functional connections, involving midbrain and thalamic areas as well as sensory and motor cortex, were strongly correlated to the performance (**Fig. 7C, right**). Consequently, when we averaged connections based on their link between cortical and subcortical ROIs (cortico-cortical, cortico-subcortical, and subcortico-subcortical), we found a distinct pattern of connections that showed strong correlation to task performance in each block type (p < 0.01, F_(2,15000)_ = 68.1, interaction of area category by block type, 2-way ANOVA) (**Fig. 7G**). Subsequent post-hoc analysis revealed that the cortico-cortical and cortico-subcortical behavior-ΔFC correlations were higher in the novel blocks compared to familiar blocks (p < 0.01, Tukey-Kramer test), while the relationship between subcortico-subcortical ΔFC and behavioral performance was lower in novel blocks and higher in familiar blocks (p < 0.01) (**Fig. 7G**).

We performed a similar analysis between ΔFC and RT across novel and familiar blocks (**Fig. 7B and D**). Here we observed a negative correlation between behavior and ΔFC in cortical areas but a positive correlation between behavior and ΔFC in midbrain and thalamic connections in both novel and familiar blocks (**Fig. 7F and H**). Although there was a significant interaction of area category by block type (p < 0.01, F_(2,15000)_ = 4.6) there was no significant difference in subcortico-subcortical connections (p = 1.0, Tukey-Kramer test). This suggests that RT was associated with subcortical FC in a similar manner in both blocks. Interestingly, the pattern of correlation between RT and ΔFC was similar to that with task performance in familiar blocks (**Fig. 7C and D**). This result suggests that the brain-wide networks associated with learning novel associations that are modulated by dopaminergic antagonists are largely separable from those associated with memory-based choices to familiar stimuli or response times. We also conducted the same analysis with behavioral data normalized for each subject (z-transformed). The networks correlated to each behavior matched those shown in **Figure 7**; cortico-cortical and cortico-subcortical behavior-ΔFC correlations were higher in the novel blocks compared to familiar blocks and subcortico-subcortical behavior-ΔFC correlations were lower in novel blocks than that in familiar blocks (p < 0.01, Tukey-Kramer test). No significant difference in subcortico-subcortical connections was observed in the relationship between the RT and ΔFC (p = 0.40).

Finally, we performed a correlation analysis between ΔFC and RL model parameters that were computed by fitting the animals’ choice data in novel blocks with a standard two-parameter RL model (**Fig. 3**). Because our model fitting analysis showed a selective change in inverse temperature following haloperidol, we expected to observe a stronger correlation between ΔFC and inverse temperature than that between ΔFC and learning rate. As predicted, ΔFC showed a strong and negative correlation to inverse temperature (> 95% confidence interval), while their correlation to learning rate was less pronounced (**Fig. 8A-C**). Strong correlations were observed in cortico-cortical and cortico-subcortical connections preferentially with inverse temperature (**Fig. 8D,** p < 0.01, F_(2,15000)_ = 46.4, interaction of area category by RL parameter, 2-way ANOVA), which mirrored the pattern observed when we analyzed correlation between ΔFC and performance in novel blocks (**Fig. 7G**), suggesting an overlap of the circuits associated with the degree of exploration and learning performance.

**Figure 8.**
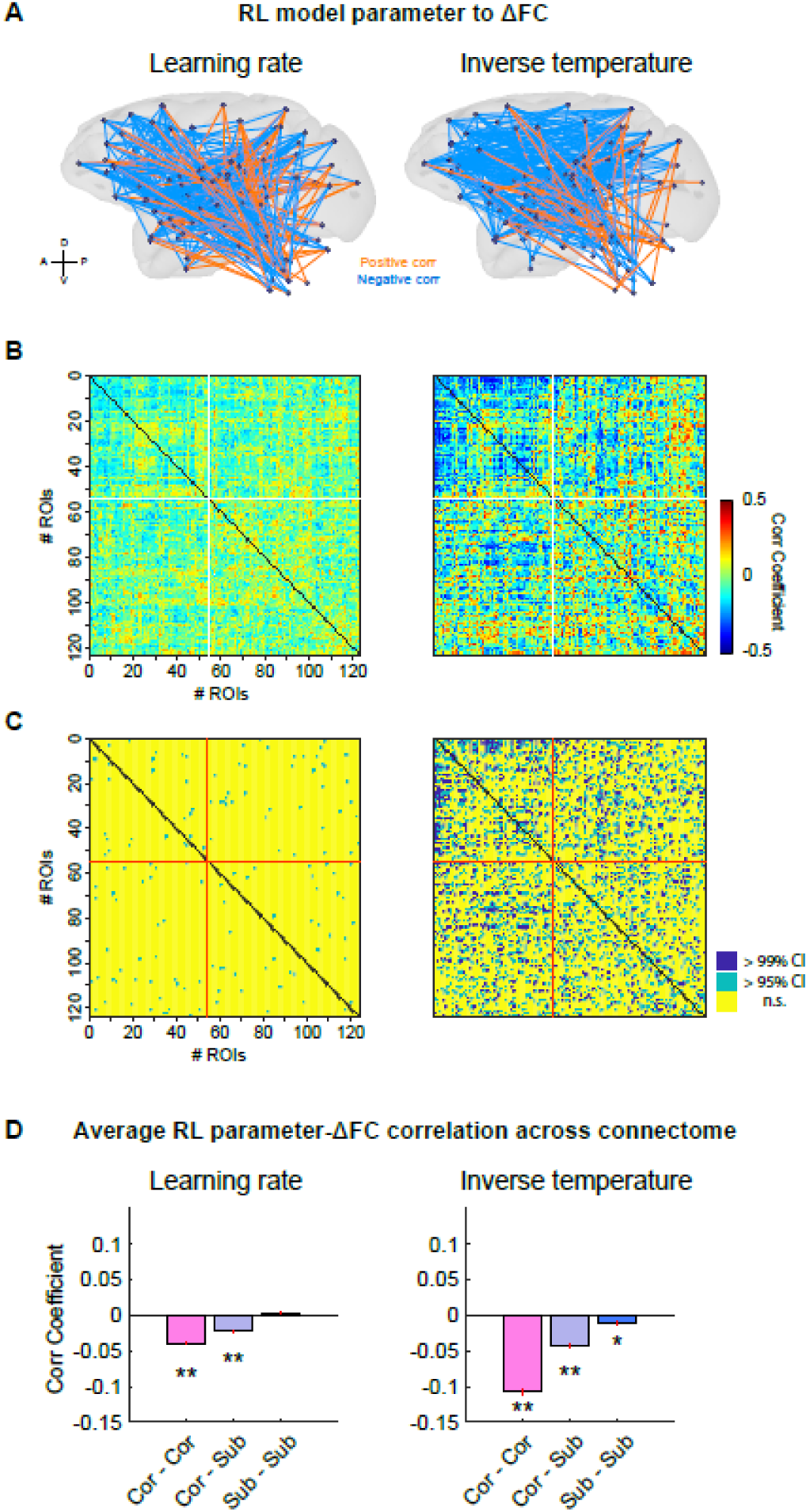
Whole-brain network correlation to reinforcement learning-model parameters across all drug conditions. (**A**) Strength of RL model parameters-ΔFC correlation projected into a brain map (Left: learning rate, right: inverse temperature). The strength and direction of changes in FC (top 5% of connections) are depicted as the width and color of lines (orange: positive, blue: negative) respectively. (**B**) Correlation matrix depicting the relationship between changes in FC and RL model parameters. Colors indicate RL model parameter-ΔFC correlation coefficient. (**C**) Functional connections (ROI pairs) that showed a significant correlation between FC changes and RL parameters (> 95% CI, permutation test) are shown in the matrix used in **B**. (**D**) Bar plots depicting average RL model parameters-ΔFC correlation coefficients calculated for cortico-cortical, cortico-subcortical, and subcortico-subcortical connections separately. **p < 0.01, rank-sum test. Conventions are the same as Fig. 7.

In sum, our analysis directly correlating behavior and resting-state FC changes induced by dopaminergic receptor antagonists revealed distinct neural networks that were associated with specific behavioral domains.

## Discussion

Here we conducted concurrent behavioral and resting-state fMRI experiments in macaque monkeys to assess the impact of dopamine D1 and D2 receptor antagonists on the brain-wide networks that support learning and motivation. Administration of the D1 receptor antagonist SCH-23390 reduced performance on a probabilistic learning task and reduced resting-state FC in cortico-cortical and fronto-striatal networks. By contrast, administration of the D2 receptor antagonist haloperidol improved performance on the same task and increased FC in cortical networks. When we looked for relationships between behavior and changes in FC induced by D1/D2 antagonists, we found that effects of dopaminergic manipulation related to learning were associated with cortico-cortical connections, whereas the effect on motivational aspects of task performance were associated with subcortical FC. Taken together, our results identified distinct brain-wide networks that underlie the impact of D1 and D2 antagonists on learning and motivation.

### The role of D1 and D2 receptors in learning and memory-based choices

The effects of DA receptor manipulation on behavior have been extensively studied in both humans and animals. Past reports using rats or macaques showed that the administration of D1 antagonist SCH-23390 and D2 antagonists raclopride or haloperidol induced opposing effects in reward-based learning and probabilistic choices (Sawaguchi and Goldman-Rakic, 1991; Eyny and Horvitz, 2003; Zeeb et al., 2009; St Onge et al., 2011; Puig and Miller, 2012; Hori et al., 2021; Jenni et al., 2021). Interestingly, unlike the robust behavioral effects observed in past studies using animal subjects, relatively mixed effects of D2 antagonism were reported in the studies using healthy humans as subjects. For instance, several studies reported that D2 antagonism enhanced reward-related signals in healthy human subjects (Jocham et al., 2011; Kahnt et al., 2015; Clos et al., 2019). In contrast, other studies reported that D2 antagonists lacked a clear effect on exploration/exploitation behaviors in a reinforcement learning task (Chakroun et al., 2020) or even impaired reinforcement learning by disrupting reward prediction error signaling (Pessiglione et al., 2006; Eisenegger et al., 2014; Diederen et al., 2017). These differences could be derived from individual variability in baseline dopamine levels (Cools and D’Esposito, 2011) and the choice of the dose given to participants (Chakroun et al., 2020), or due to dose-dependent difference in the main site of action of haloperidol (i.e., pre-synaptic vs. post-synaptic effects), as we discuss later. In addition, there is a possibility that the difference in task design across studies could lead to such a discrepancy in the drug’s effect on the overall choice performance. In the human studies that observed deficits in performance following haloperidol treatment, subjects performed two-option probabilistic tasks (Pessiglione et al., 2006; Eisenegger et al., 2014). By contrast, in the current study subjects chose between three stimuli that were probabilistically rewarded in each novel block, which likely made value-based learning harder and favored more prolonged exploration. Thus, it is possible that increasing the degree of exploration was advantageous in our task but was actually disadvantageous in the two-option tasks. Indeed, fitting a two-parameter reinforcement learning model to the subjects’ choices showed that haloperidol selectively decreased the inverse temperature parameter in novel blocks. Notably, this change in the degree of exploration was consistent with the above human studies even though the effect on correct performance was the opposite. This highlights that the haloperidol dose that we used here did not simply change subjects’ performance via modulating motivation or attention, but specifically impacted their behavioral strategies including the degree of exploration. Additionally, our task design tested animals in both novel and familiar conditions, allowing us to dissociate the behavioral effects of drugs on learning from those on motor or motivational functions.

Our behavioral results were overall consistent with the existing literature; D1 antagonist SCH impaired and D2 antagonist haloperidol facilitated the performance of our monkeys in novel blocks (**Fig. 2**). Notably, DA receptor manipulation in this range did not affect the performance in the familiar block, suggesting that the actions of DA through D1 and D2 receptors play a specific role in new association learning rather than choices in general. In addition to the effects on learning performance, we also observed a change in subjects’ RTs specifically in the SCH sessions. Notably the impact of SCH on RT was observed in both novel and familiar blocks, suggesting that the effect of DA receptor manipulation on motivation or motor function is dissociable from the effects on learning. Our model fitting analysis further revealed a selective and dose-dependent decrease in the inverse temperature parameter following administration of the D2 receptor antagonist haloperidol, suggesting that this improved the animals’ performance by slightly increasing the level of exploration. The negative effect of D2 antagonism on the inverse temperature parameter without appreciably impacting the learning rate is consistent to previous findings in human subjects (Pessiglione et al., 2006; Eisenegger et al., 2014). Taken together, our behavioral analyses demonstrated contrasting behavioral effects following systemic manipulation of D1 and D2 receptors, where D2 receptor antagonism specifically impacted choice consistency during learning.

It is important to note, however, that the effect of haloperidol administration on behavior could be interpreted as being predominantly caused by its affinity for pre-synaptic D2 receptors on striatal neurons. On this view, haloperidol at low doses could inhibit pre-synaptic D2 receptors, which is thought to lead to increased DA release from the axon terminal. If this was the case, the effects of haloperidol administration in our experiments would be the result of increased DA release as opposed to haloperidol antagonistically acting on post-synaptic D2 receptors. Indeed, the doses we used in the current study were lower than the doses typically used in human studies or in clinical settings where more than 1-2 mg haloperidol (equivalent to 14-28 ug/kg for a 70 kg male subject) was used (Pessiglione et al., 2006; Chakroun et al., 2020). There are several reasons why we believe that this is unlikely to be the case. First, our haloperidol dosage was determined based on a prior PET study using drug-naïve macaques, where single administration of 10 ug/kg haloperidol occupied 80% of striatal D2 receptors (Hori et al., 2021). By contrast, in healthy humans, single administration of 3 mg (42 ug/kg for a 70 kg male) haloperidol occupies only 35-65% of D2 receptors in the striatum (Ishiwata et al., 2006; Lim et al., 2013). Notably, daily treatment with 3 mg haloperidol leads to 80% D2 occupancy after several days in humans (Zipursky et al., 2005; Lako et al., 2013; Lim et al., 2013). In addition to this, there appear to be differences between effective doses of haloperidol across species that must be considered when comparing studies of humans and animal models (Kapur et al., 2000; Mukherjee et al., 2001). Therefore, it is likely that our haloperidol doses were not low in terms of D2 receptor occupancy level, and that their administration to drug-naïve macaques sufficiently induced post-synaptic effects that are equivalent to the previous human studies. We acknowledge, however, that without further investigation with higher doses of haloperidol, and/or additional investigation using dopamine agonists, we cannot rule out the possibility that our haloperidol results are at least partially accounted for by its action to pre-synaptic D2 receptors. Future study should delineate among these possibilities by testing both agonists and antagonists in wider dose ranges.

### Dopaminergic modulation of fMRI resting-state functional connectivity

Previous studies have mainly analyzed neural effects of DA receptor manipulation by focusing on specific areas such as prefrontal cortex and striatum (Wang et al., 2004; Noudoost and Moore, 2011; Puig and Miller, 2012; Yael et al., 2013; Puig and Miller, 2015; Kunimatsu and Tanaka, 2016). One advantage of our resting-state fMRI approach is that it can identify drug effects on intrinsic networks free from the indirect impact of drug-induced behavioral changes. Further, our neuroimaging protocol uses a low level of anesthesia to preserve resting-state FC in macaque monkeys meaning that brain-wide FC patterns are still sensitive to pharmacological treatment (Fujimoto et al., 2022; Elorette et al., 2024). Using this approach, our whole-brain connectome analyses revealed contrasting effects of SCH and haloperidol, particularly in cortico-cortical and cortico-subcortical connections, mirroring the changes in learning performance induced by the same drugs (**Fig. 5**). In addition to the known effects on fronto-striatal circuits and fronto-parietal networks, diverse cortical regions including temporal areas and thalamic nuclei were involved. The present results highlight that large-scale functional networks are recruited by DA receptor modulation to influence various cognitive and motor functions.

Interestingly, the pattern of effects on functional connectivity after D2 receptor manipulation did not simply reflect the known distribution of this receptor subtype within the primate brain, which is mainly localized to the striatum (Suhara et al., 1999; Tsukada et al., 2005; Froudist-Walsh et al., 2021; Hori et al., 2021). It is unlikely that the non-specific binding of haloperidol to D1 receptors caused changes in cortical areas, as the overall direction of the effects was the opposite between those drug conditions. One possibility is that the haloperidol induced substantial neural changes through interactions with D2 receptors expressed in cortical neurons, including presynaptic autoreceptors (Beaulieu and Gainetdinov, 2011; Cools and D’Esposito, 2011). Indeed, previous studies demonstrated that cortical D2 receptors are functionally relevant (Narendran et al., 2009; Narendran et al., 2014) and associated with positive symptoms in schizophrenia (Suhara et al., 2002; Mizrahi et al., 2007), although the profile of cortical D2 receptors is still unclear due to technical limitations (Tritsch and Sabatini, 2012). This question could be addressed by recording neuronal activity from D1 and D2 receptor expressing neurons in both cortical and striatal regions.

### Dissociable neural networks for distinct dopamine-dependent behaviors

Past studies have demonstrated that resting-state FC can be used to predict the behavioral effects of pharmacological treatments on learning, memory recall, and attention, in both humans and macaques (Li et al., 2013; Kohno et al., 2014; Fujimoto et al., 2022). Our within-subject behavior-connectivity correlation analysis revealed distinct brain networks where connectivity was correlated with task performance or RT (**Fig. 7**). The network that we identified related to learning performance included fronto-striatal and fronto-parietal circuits and largely overlaps with networks known to be more active when subjects are learning reward-based associations (Cools et al., 2004; Cohen, 2008; Chadick and Gazzaley, 2011; Frank and Badre, 2012; Sescousse et al., 2013; Gilmore et al., 2015). Further correlation analysis between the connectome and RL parameters revealed that these brain networks are associated with variation in the inverse temperature, suggesting that the dopamine receptor manipulation predominantly affects the degree of exploration rather than the rate of value updating. It is noteworthy that the network reflecting task performance in familiar blocks, including midbrain and thalamic nuclei, largely overlaps with the network of brain areas correlated with RT. That different behavioral domains engaged the same network of areas indicates that this system may play a central role in motivation or motor control of executing a choice after learning has occurred. Indeed, a recent study demonstrated that silencing of the ventral tegmental area to ventral striatum pathway in macaques affected motivation but did not impair reinforcement learning (Vancraeyenest et al., 2020). Thus, our analysis revealed distinct neural networks where dopamine takes action to modulate behaviors in primates.

### Conclusion

Dopaminergic signaling, especially an optimal balance between D1 and D2 receptor-dependent modulation, is critical for normal learning (Seeman, 1987; Takahashi et al., 2012), and its alteration may contribute to the basis of schizophrenia (Sedvall and Karlsson, 1999; Yun et al., 2023). The similarity of the dopaminergic system between non-human primates and humans (Berger et al., 1991; Raghanti et al., 2008) means that our findings have implications for the brain-wide actions of antipsychotics in humans. Thus, our data provide evidence that the cognitive effects of D1/D2 receptor modulation are related to altered functional connections among cortical areas and reveal a possible mechanism through which systemic pharmacological DA receptor manipulation contributes to ameliorating aberrant cognition.

## Acknowledgments

A.F., C.E., B.E.R., and P.H.R. are supported by grants from the BRAIN initiative (R01MH117040). B.E.R. is supported by grants from NIMH (R01MH111439) and NINDS (R01NS109498). A.F. is supported by Overseas Research Fellowship from Takeda Science Foundation and a Brain & Behavior Research Foundation Young Investigator grant (#28979). We would like to thank Dr. Paula Croxson for providing the foundation on which this work was built, Dr. Yukiko Hori for advising on drug preparation, and Jairo Munoz and Niranjana Bienkowska for assistance with data acquisition. For help with fMRI data pre-processing and analysis we thank Drs Paul Taylor and Alex Franco, respectively. We also thank Drs. Jacqueline-Marie Ferland, Dan Iosifescu, and Takafumi Minamimoto for their comments on the earlier version of the manuscript.

## Conflict of interest

The authors declare no competing financial interest.

## Author contributions

A.F. designed the study. A.F. and C.E. collected the behavioral data. A.F., C.E., S.H.F., and L.F. collected the imaging data. A.F. analyzed the data. A.F., C.E., P.H.R., and B.E.R. wrote the original draft. All authors edited the paper.

## Data Availability

The data that support the findings of this study are available from the corresponding authors upon reasonable request.

## Notes

### Competing Interest Statement

The authors have declared no competing interest.

### Summary of Updates

Figures 1, 2, 4, 5, 6, and 7 revised; Figures 3 and 8 added. Text revised.

